# Strengthening connectivity between premotor and motor cortex increases inter-areal communication in the human brain

**DOI:** 10.1101/2023.02.15.528606

**Authors:** Jelena Trajkovic, Vincenzo Romei, Matthew F. S. Rushworth, Alejandra Sel

## Abstract

The ventral premotor cortex (PMv) is an important component of cortico-cortical pathways mediating prefrontal control over primary motor cortex (M1) function. Paired associative stimulation (ccPAS) is known to change PMv influence over M1 in humans, which manifests differently depending on the behavioural context. Here we show that these changes in influence are functionally linked to PMv-M1 phase synchrony changes induced by repeated paired stimulation of the two areas. PMv-to-M1 ccPAS leads to increased phase synchrony in alpha and beta bands while reversed order M1-to-PMv ccPAS leads to decreased theta phase synchrony. These changes are visible at rest but they are predictive of changes in oscillatory power in the same frequencies during movement execution and inhibition, respectively. The results unveil a link between the physiology of the motor network and the resonant frequencies mediating its interactions, and provide a putative mechanism underpinning the relationship between synaptic efficacy and brain oscillations.

## Introduction

The synchronisation of neuronal oscillations is increasingly being appreciated as a key element in communication between brain areas (Fries, 2005, Salinas and Sejnowski, 2001, Fries, 2015). This is because the oscillatory properties of the neurons may determine their ability to send and receive electrical signals. Interregional cortico-cortical coupling, for example, only occurs when oscillations in different neuronal sets create rhythmic opportunities during which cells simultaneously increase or decrease their readiness to transfer information, regulating the timing of action potentials in a circuit-dependent manner (Fries, 2015, Fries, 2005, Salinas and Sejnowski, 2001). Importantly, it is thought that it is only possible for different sets of neurons to oscillate synchronously when they share a common physiological substrate (van Ede et al., 2018, Wang, 2010). Here we test this idea by carrying two types of reversible manipulations of a human brain circuit that have been established to either transiently increase or decrease physiological interconnectedness. We then examine whether the two types of intervention result in increased or decreased oscillatory coupling between activity in the component areas of the circuit.

The neural circuit mediating action control that runs between the ventral premotor cortex (PMv) and primary motor cortex (M1) is the one in which we conduct our experiments. Executive control processes in prefrontal cortical areas can only control, sculpt, or influence sensory and motor processes if there are anatomical connections between prefrontal cortex and sensory and motor areas. One of the most direct cortico-cortical routes through which prefrontal cortex may influence the motor system is via PMv. First, there is a substantial and monosynaptic projection from prefrontal cortex to PMv (Dum and Strick, 2005). Moreover, in turn it is well established that PMv exerts a powerful influence over M1 and that changes in PMv-M1 connectivity are functionally relevant and correlated with motor control both in humans and macaques (Shimazu et al., 2004, Cerri et al., 2003, Prabhu et al., 2009b, Fiori et al., 2018, Davare et al., 2008, Davare et al., 2009, Davare et al., 2010). Although the projections from premotor areas, including PMv, to M1 are monosynaptic and excitatory, many are made onto inhibitory interneurons(Tokuno and Nambu, 2000), ensuring that PMv is able to exert both a facilitatory influence over M1 during action execution as well as an inhibitory influence at rest and when an action needs to be stopped (Shimazu et al., 2004, Prabhu et al., 2009a, Davare et al., 2008, Davare et al., 2009, Davare et al., 2010, Buch et al., 2010). Therefore, the influence exerted by PMv on M1 is state dependent, being mostly facilitatory during action initiation but inhibitory when actions are to be curtailed or stopped (Neubert et al., 2010, Buch et al., 2010, Davare et al., 2008, Buch et al., 2011). At the same time, initiation and cessation of movements have been linked, respectively, to decreases and increases in beta and theta frequency oscillations (Zhang et al., 2008, Picazio et al., 2014, Tsujimoto et al., 2010, Harper et al., 2014). These may operate as spectral fingerprints of top-down executive control (Tsujimoto et al., 2010, Tsujimoto et al., 2006, Yamanaka and Yamamoto, 2010, Harper et al., 2014, Helfrich et al., 2018, Helfrich et al., 2019). Similarly, interregional alpha coupling is argued to reflect information transfer from PMv, and adjacent prefrontal areas, to M1 (Hughes et al., 2018, Liebrand et al., 2018).

The causal influence exerted by PMv over M1 can be studied by stimulating PMv with a single pulse of transcranial magnetic stimulation (TMS) shortly (6-8ms) before stimulating M1 with another TMS pulse, making it possible to examine how M1 activity evolves as directly influenced by PMv (Neubert et al., 2010, Buch et al., 2010, Davare et al., 2008, Davare et al., 2009, Davare et al., 2011, Casarotto et al., 2022). Although the impact of the first pulse in PMv is spatially circumscribed (Romero et al., 2019), it alters the activity in PMv neurons that project to M1 (Shimazu et al., 2004, Cerri et al., 2003, Prabhu et al., 2009b). However, when such a paired stimulation protocol is applied in a repetitive manner, it is possible to strengthen the influence that PMv exerts over M1 (Buch et al., 2011, Johnen et al., 2015, Chiappini et al., 2020, Sel et al., 2021). Such a procedure is referred to as cortico-cortical paired associative stimulation (ccPAS) (Chiappini et al., 2022, Turrini et al., 2022, Chiappini et al., 2018, Romei et al., 2016, Sel et al., 2021, Buch et al., 2011, Johnen et al., 2015). The evoked effects have been described as Hebbian in nature (Huang et al., 2017, Koch et al., 2013) and are thought to tap into mechanisms of spike-timing dependent plasticity (STDP) (Buch et al., 2011, Johnen et al., 2015). On the other hand, when the ccPAS is applied to the same cortical areas (PMv and M1) but with a longer inter-pulse interval (IPI) or in the reversed temporal order (M1-to-PMv ccPAS), then the evoked effects have been described as anti-Hebbian (Koch et al., 2013), and result in a diminished influence of PMv on M1 (Buch et al., 2011, Johnen et al., 2015, Sel et al., 2021). These effects have been established by measuring changes in M1 motor related activity (Johnen et al., 2015, Buch et al., 2011) and the coupling of blood oxygen level dependent (BOLD) signals in PMv and M1 before and after ccPAS (Johnen et al., 2015).

From these observations, it is evident that changes in synaptic efficacy and connectivity strength in the PMv-M1 pathway are functionally significant and related to top-down motor control. The impact of ccPAS on PMv-M1 connectivity strength can also be visualised by measuring task-related oscillatory changes (Sel et al., 2021). When examining changes in the task-related oscillations recorded prior to and after PMv-to-M1 ccPAS, frequency-specific changes in the beta band increase in contexts, such as movement production, in which M1 normally receives an excitatory influence from PMv. Such frequency specific changes are, however, context-dependent and in other settings like action cancellation, where PMv inhibits M1, it is the theta activity instead that is augmented by ccPAS (Sel et al., 2021). Taken together, the physiological properties of the PMv-M1 pathway and its resonant dynamics, in tandem with the powerful capacity of ccPAS to selectively manipulate the strength of connections between human cortical areas, make them an ideal test bed for hypothesis about how neuronal coupling reflect brain circuitry and function. However, the degree to which various patterns of frequency-specific change are dependent on the current behavioural state (moving or stopping), or indicative of fundamental features of PMv-M1 pathway anatomy and oscillatory architecture is still largely unknown. To date, effects in each frequency band have been recorded in specific behavioural states. But it is possible that PMv-to-M1 ccPAS effects are more prominent at higher frequencies such as alpha and beta bands while reversed order M1-to-PMv ccPAS effects are more prominent at lower frequencies such as theta. Testing this possibility requires, first, analysis of ccPAS effects in the absence a motor task, for example during rest and, second, measures of inter-regional phase synchrony as opposed simply to changes in power in various frequency bands averaged across brain regions (Sel et al., 2021).

Here we aimed to test the impact of either increasing or decreasing the strength of connections between PMv and M1 on interregional coupling of neural responses by measuring EEG activity from prefrontal and motor cortical areas at rest in two blocks (referred to as Baseline and Expression blocks) before and after ccPAS. In two participant groups, we used two patterns of magnetic stimulation that either increased or decreased connectivity strength between PMv and M1. In participant Group 1 we applied 15 minutes of ccPAS in which each TMS pulse over PMv was followed by a TMS pulse over M1 after a 6 or 8 ms IPI (PMv-to-M1-ccPAS). Before and after ccPAS, we recorded EEG cortical activity at rest. Moreover, in participant Group 2, we reversed the order of ccPAS stimulation i.e., applying the first TMS pulse in each pair over M1 and the second over PMv (Fig. 1) to investigate whether changes in oscillatory coupling were dependent on ccPAS stimulation order. Importantly, comparison of the two protocols allows us to control for the effects of inducing activity in each individual area as opposed to carrying out a manipulation of the pathway interconnecting them; an identical number of pulses were applied both to PMv and M1 in both participant groups 1 and 2 but the two protocols should have opposite effects on PMv to M1connection strength (Buch et al., 2011). We hypothesised that augmenting the strength of PMv-M1 connections, by stimulating the PMv-to-M1 pathway, may result in increased interregional phase synchrony between pre-motor and primary motor cortices; furthermore, we also predicted that diminishing PMv-M1 connectivity strength, by stimulating the M1-to-PMv pathway, may lead to phase synchrony decreases across pre-motor and primary motor regions. In addition, we hypothesised that increases and decreases in interregional coupling between PMv and M1 areas may also relate to frequency amplitude changes that are related to top-down motor control during movement control.

**Figure 1:**
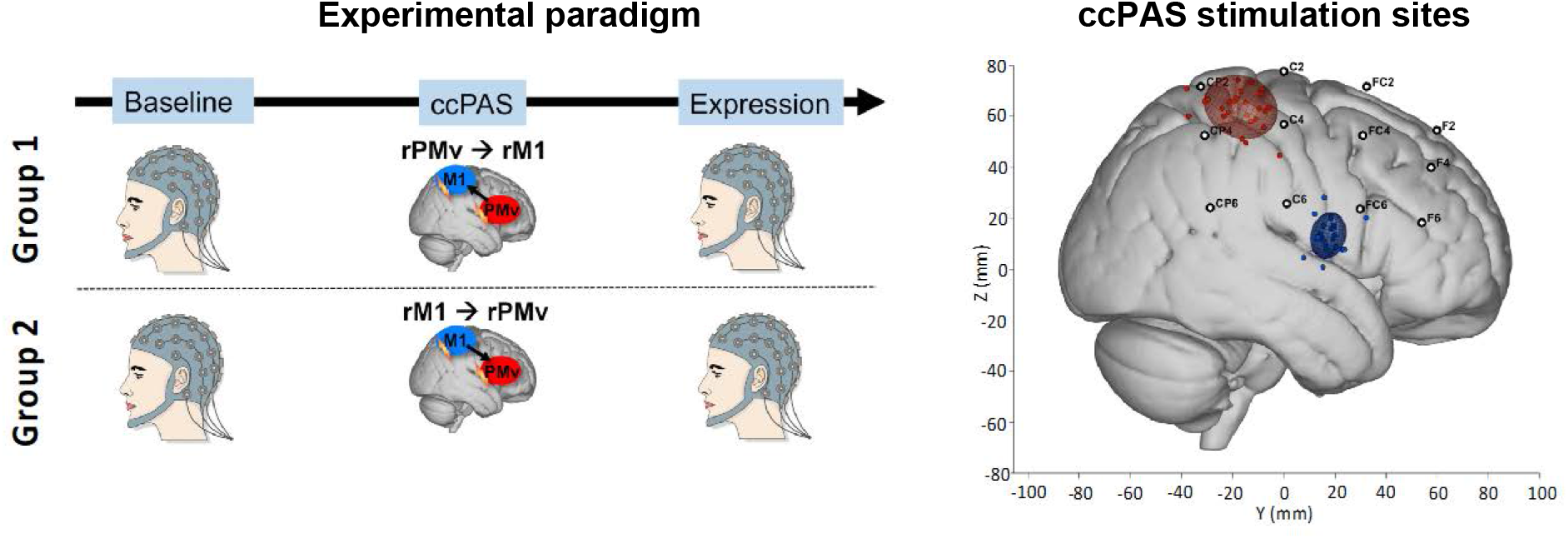
Representation of the set up for Groups 1 and 2, and individual scalp hotspot for right primary motor cortex (M1) and right ventral premotor cortex (PMv). The filled circles in red (PMv) and blue (M1) represent subject scalp hotspot and the ellipses represent the 95% group confidence for the PMV and M1 location for Group 1 and Group 2 in standardised MNI space. The right-hand panel also shows a representation of the electrode labels and locations over the stimulated areas in the right hemisphere.

## Material and methods

### Participants

36 healthy, right-handed adults participated across the two experimental groups: 18 in participant group 1 (mean age ± SD 23.61 ± 4.50; 10 female participants; 0.81 ± 0.16 handiness mean ± SD; as measured by the Edinburgh handedness inventory-adapted from (Oldfield, 1971)); 18 in participant group 2 (age: 23.05 ± 2.83; 5 female participants; 0.93 ± 0.13 handedness). All participants had no personal or familial history of neurological or psychiatric disease, were right-handed (except for one participant – handedness score .045), gave written informed consent (Medical Science Interdivisional Research Ethics Committee, Oxford RECC, No. R29477/RE004), were screened for adverse reactions to TMS and risk factors by means of a safety questionnaire, and received monetary compensation for their participation. Participants underwent high-resolution, T1-weighted structural MRI scans. Sample sizes were determined based on previous studies that have used the same ccPAS protocol to measure the influence of PMv over M1 cortical excitability (Johnen et al., 2015, Buch et al., 2011), and studies that have used EEG to investigate cortico-cortical coherence in the human brain.

### Experimental design

Both experiments started with a Baseline block, followed by a ccPAS period, and an Expression block (Fig. 1). During Baseline and Expression blocks, we recorded resting EEG data while participants fixated on a white cross on the centre of the computer screen during 3 periods of 1-minute length each, i.e., total time 3 minutes. We have previously reported the impact of the TMS protocol on a Go/No-go task (Sel et al., 2021). Participants were seated approximately 50cm from the screen in a sound and electrically shielded booth. In the two experimental conditions, the ccPAS period that intervened between Baseline and Expression blocks consisted of 15 min of ccPAS over PMv and M1 applied at 0.1 Hz (90 total stimulus pairings) with an IPI of either 6 or 8 ms. Both resting-state and task-state interactions between M1 and PMv, and adjacent areas, emerge at 6–8 ms intervals (Davare et al., 2008, Davare et al., 2009, Johnen et al., 2015, Buch et al., 2011). Precise inter pulse timing is critical if both PMv and M1 pulses are to produce coincident influences on M1 neurons. Therefore, we employed an IPI of 8 ms when testing half of the participants in session 1 and in session 2, and an IPI of 6ms in the other half of participants in each experimental session. However, because subsequent analyses that included consideration of IPI found no effect of the 2ms difference, we do not consider this difference in IPI further. In participant Group 1, the pulse applied to PMv always preceded the pulse over M1, while the opposite was true in Group 2, where M1 TMS preceded PMv TMS.

### Cortico-cortical Paired Associative Stimulation

ccPAS was applied using two Magstim 200 stimulators, each connected to 50 mm figure-eight shaped coils. The M1 “scalp hotspot” was the scalp location where the TMS stimulation evoked the largest motor evoked potential (MEP) amplitude in left first dorsal interosseous (FDI) muscle. This scalp location was projected onto high-resolution, T1-weighted MRIs of each volunteer’s brain using frameless stereotactic neuronavigation (Brainsight; Rogue Research). In contrast to the scalp hotspot, the right M1 “cortical hotspot” was the mean location in the cortex where the stimulation reached the brain for all participants in Montreal Neurological Institute (MNI) coordinates (X = 41.03 ± 6.59, Y = – 16.74 ± 9.35, Z = 63.69 ± 8.20; Fig. 1 – cortical coordinates computed using Brainsight stereotactic neuronavigation for each participant; mean cortical coordinates computed by averaging all individual’s cortical coordinates). These coordinates were similar to those reported previously (Davare et al., 2009, Buch et al., 2011, Buch et al., 2010, Johnen et al., 2015). The PMv coil location was determined anatomically as follows. A marker was placed on each individual’s MRI and adjusted with respect to individual sulcal landmarks to a location immediately anterior to the inferior precentral sulcus. The mean MNI cerebral location of the PMv stimulation was at (X = 59.66 ± 3.41, Y = 17.07 ± 6.28, Z = 14.85 ± 8.50; Fig 1) and lies within the region defined previously as human PMv (rostral part) and the adjacent inferior frontal gyrus (posterior/mid part) (Mayka et al., 2006), more precisely over areas 44d and 44v of the pars opercularis within the inferior frontal gyrus (Neubert et al., 2014), which resembles parts of macaque PMv in cytoarchitecture and connections (Croxson et al., 2005).

Resting motor threshold (RMT) of the right M1 (mean ± SD, 43.13 ± 7.22% stimulator output) was determined as described previously (Rossini et al., 1994). As in previous ccPAS studies (Neubert et al., 2010, Buch et al., 2011, Buch et al., 2010, Johnen et al., 2015), PMv TMS was proportional to RMT - 110% (47.76 ± 7.35). M1 stimulation intensity during experiments was set to elicit single-pulse MEPs of ±1 mV (47.23 ± 7.58% stimulator output). TMS coils were positioned tangential to the skull, with the M1 coil angled at ~45° (handle pointing posteriorly), and the PMv coil at ~0° relative to the midline (handle pointing anteriorly). The PMv coil was fixed in place with an adjustable metal arm and monitored throughout the experiment. The M1 coil was held by the experimenter. Left FDI electromyography (EMG) activity was recorded with bipolar surface Ag-AgCl electrode montages. Responses were bandpass filtered between 10 and 1000 Hz, with additional hardwired 50 Hz notch filtering (CED Humbug), sampled at 5000 Hz, and recorded using a CED D440-4 amplifier, a CED micro1401 Mk.II A/D converter, and PC running Spike2 (Cambridge Electronic Design).

### EEG recording and analysis

EEG was recorded with sintered Ag/AgCl electrodes from 64 scalp electrodes mounted equidistantly on an elastic electrode cap (64Ch-Standard-BrainCap for TMS with Multitrodes; EasyCap). All electrodes were referenced to the right mastoid and re-referenced to the average reference off-line. Continuous EEG was recorded using NuAmps digital amplifiers (Neuroscan, El Paso, Texas 1000Hz sampling rate).

Off-line EEG analysis was performed using Fieldtrip (Oostenveld et al., 2011) and the connectivity analysis was done using EEGLAB (EEGLAB v2021.1, University of San Diego, San Diego, CA). First, the data was down-sampled to 500Hz and digitally band-pass-filtered between 1-40 Hz. Bad/missing channels were restored using a FieldTrip based spline interpolation. Next, the data were segmented into 2s intervals, which resulted in a total of 90 segments recorded before and 90 segments recorded after the ccPAS. Automatic artefact rejection was combined with visual inspection for all participants eliminating large technical and movement related artefacts. Physiological artefacts such as eye blinks and saccades were corrected by means of independent component analysis (RUNICA, logistic Infomax algorithm) as implemented in the FieldTrip toolbox. Those independent components (most often one or two) whose timing and topography resembled the characteristics of the physiological artefacts were removed. The signal was re-referenced to the arithmetic average of all electrodes.

### Connectivity analysis

Phase connectivity was estimated in the sensor space, via weighted phase-lag index (wPLI) (Vinck et al., 2011). This is a measure of phase lag-based connectivity, that accounts for non-zero phase lag/lead relations between two time series signals, Therefore, it is a measure insensitive to volume conduction and noise of a different origin, considered optimal for exploratory analysis as it minimizes type-I errors (Cohen, 2014, Cohen, 2015).

In order to obtain wPLI values, time series data was first transformed into the time-frequency domain via convolution with a family of complex Morlet wavelets (number of cycles increased from 5 to 18 in logarithmic steps). Therefore, for frequencies ranging from 3 to 30 Hz in 1-Hz steps, first convolution by frequency-domain multiplication was performed and then the inverse Fourier transformation was taken. Phase was defined as the angle relative to the positive real axis, and phase differences were then computed between all possible pairs of electrodes. Finally, wPLI was calculated as the absolute value of the average sign of phase angle differences, whereas vectors closer to the real axis were de-weighted.

### Statistical analysis

Once wPLI values were extracted for every frequency bin (3 to 30 Hz in 1 Hz steps) and epoch, they were averaged to obtain distinct values for each block (Baseline, Expression) and each frequency band (theta: 4-7 Hz, alpha: 8-13 Hz, beta: 14-30 Hz). Subsequently, non-parametric permutation-based analysis (1000 iterations) was performed to compare the connectivity between Baseline and Expression phase, and to obtain phase connectivity difference maps of distinct ROIs: right (stimulated) hemisphere (F2, F4, F6, FC2, FC4, FC6, C2, C4, C6, CP2, CP4, CP6), left hemisphere (F1, F3, F5, FC1, FC3, FC5, C1, C3, C5, CP1, CP3, CP5) and interhemispheric connectivity between left and right ROI. Sensor differences with z-values corresponding to p<0.05 were retained as significant. The connectivity index of each condition was then estimated using the formula: CI = (sp_pos – sp_neg)/ sp_total, where sig_pos are connections that are significantly higher after ccPAS with respect to before; sig_neg those that are lower, and sp_tot are all possible connections (Alekseichuk et al., 2016).

Furthermore, another permutation test was introduced in order to calculate the significance threshold for CI. Specifically, wPLI matrices of all frequencies, ROIs and groups were randomly permuted and compared 1000 times to obtain the distribution of randomly obtained differences in wPLI. Connectivity indices that exceeded 95% confidence interval were considered statistically significant (negative threshold=-0.088; positive threshold=0.089) (Alekseichuk et al., 2016).

In addition, in order to directly contrast changes in connectivity between the groups (Group 1, Group 2), across different electrode clusters (theta, alpha and beta significant clusters) and frequencies (theta, alpha and beta frequency), a 2 x 3 x 3 repeated measures mixed-model three-way ANOVA was used.

Finally, to compare resting-state connectivity differences between Expression and Baseline blocks, obtained here in the current analysis, with the changes in time-frequency responses during the Go/No-go task, a robust skipped correlation analysis was used (Pernet et al., 2013). Specifically, the mean of normalized differences between significant sensors in the beta (significant sensors in the Group 1 analysis of PMv-to-M1 ccPAS) and theta band (significant sensors in the Group 2 analysis of reversed order PMv-to-M1 ccPAS) were computed for both groups, by using the formula: 100* (mean_post-mean_pre)/mean pre (Alekseichuk et al., 2016). These values, across both groups and in both beta and theta bands were then correlated with the differences in time-frequency responses in the same frequency band that had been found in the same participant sample during a Go/No-go task (Sel et al., 2021).

## Results

In two participant groups, Group 1 (N=18) and Group 2 (N=18), we investigated, respectively, whether increasing or decreasing coupling across motor and premotor areas led to changes in EEG oscillatory coherence at rest. We contrasted the effects of the two types of ccPAS, repeated paired stimulation of PMv followed by M1 (Group 1) or, *vice versa*, M1 followed by PMv (Group 2) on time-frequency oscillatory responses, recoded during rest. We focused on motor-relevant frequency bands theta, alpha and beta (4-30Hz) in fronto-central and centro-partietal electrodes (Methods: EEG recording and analysis) known to reflect top-down control of motor activity during rest (Zhang et al., 2008, Picazio et al., 2014, Sauseng et al., 2009, Sauseng et al., 2013, Tsujimoto et al., 2010, Tsujimoto et al., 2006, Harper et al., 2014, Yamanaka and Yamamoto, 2010). First, we examined a cluster of electrodes located around the areas of stimulation in the right hemisphere (right region of interest – right ROI). However, because the effects of ccPAS can occur across hemispheres (Neubert et al., 2010, Sel et al., 2021) we expanded our analysis to include a bilateral group of electrodes spanning homologous areas in both hemispheres (left ROI and interhemispheric ROI).

To assess the effect of ccPAS over PMv and M1 on their inter-site phase coupling, we used the weighted phase lag index (wPLI): a phase lag-based measure not affected by volume conduction and not biased by the sample size (Vinck et al., 2011, Cohen, 2014). Subsequently, non-parametric permutation analysis was used to compare wPLI before and after ccPAS stimulation between different sensors of ROIs, as well as to assess the significance of global connectivity changes within ROIs in distinct frequency bands (for details see Methods).

The results indicated that the frequency and directionality of connectivity changes are dissociable for the two ccPAS protocols. Specifically, in Group 1 (PMv-to-M1 ccPAS) there was an increase in the connectivity within the right ROI, in the alpha and beta band (CIalpha = 0,091 > CI_threshold_; CI_beta_ = 0,091 > CI_threshold_) but not in the slower theta band (CI_theta_=0,030 < CI_threshold_) (Fig. 2).

**Figure 2:**
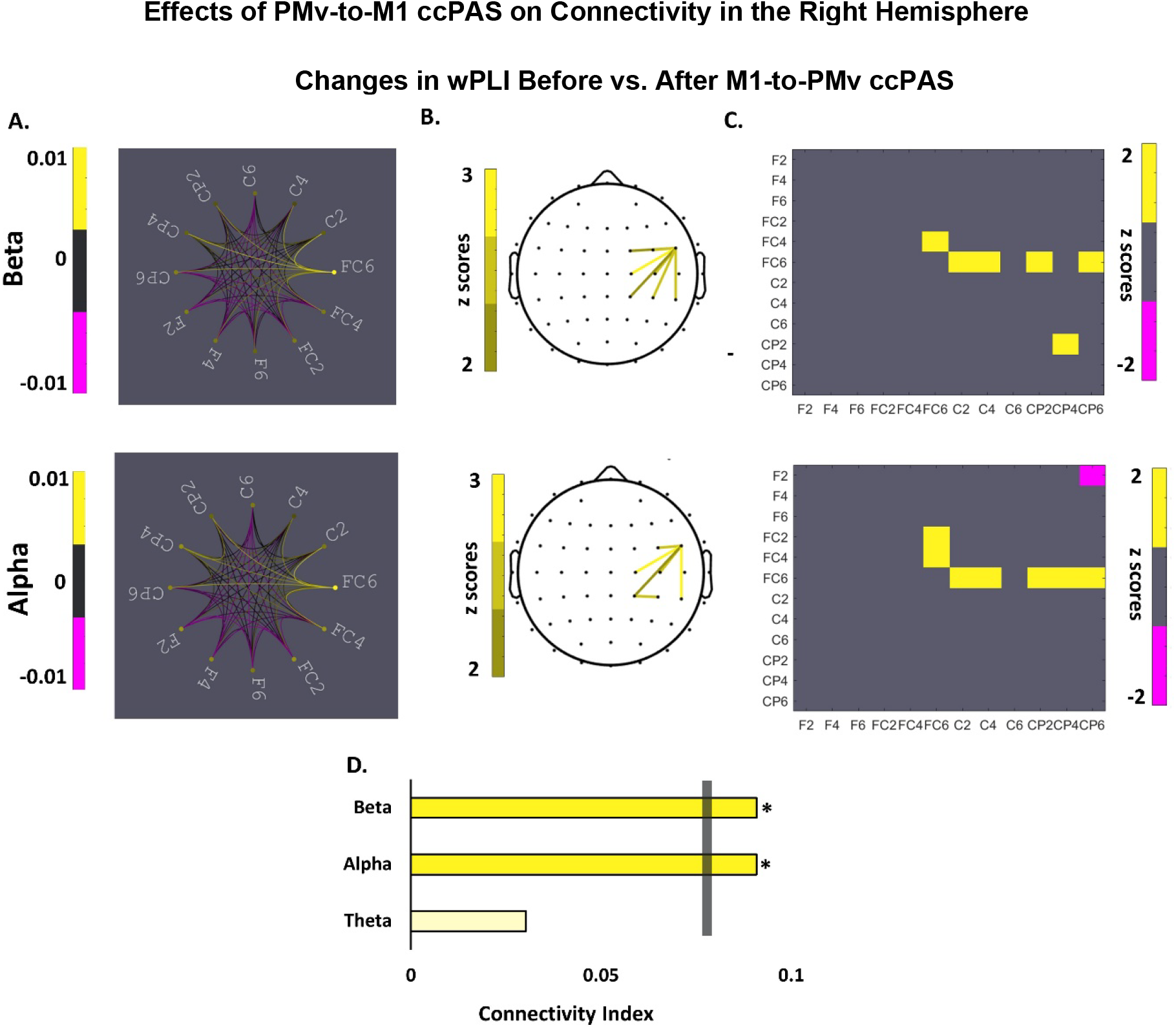
Changes in interregional coupling between PMv and M1 when contrasting activity recorded before and after PMv-to-M1 ccPAS. (A) Representation of the connectivity strength measured by the weighted phase lag index (wPLI) across all electrodes included in the regions of interest for the beta (top) and alpha (bottom) bands. (B) Topographical representations of the electrodes showing significant increased coupling in the beta (top) and alpha (bottom) bands after PMv-to-M1 ccPAS. (C) Connectivity matrix indicating significant interregional coupling increases between the electrodes of interest. (D) Representation of the connectivity index for each of the frequency bands of interest; grey vertical bar shows the statistical threshold. Yellow and pink ink indicate increases or decreases in interregional coupling after ccPAS, respectively.

Moreover, the sensor with the highest number of significant increases in connectivity with other electrodes within the ROI (electrode FC6) was the sensor closest to the area targeted by PMv TMS (see Fig. 1 & 2). On the other hand, in Group 2 (M1-to-PMv ccPAS) there was a significant decrease in connectivity within the right ROI in the theta band (CI_theta_=-0,091 > CI_threshold_) (Fig. 3), but not in the alpha and beta band (CI_alpha_ = −0,045 < CI_threshold_; CI_beta_= −0,030 < CI_threshold_). Interestingly, the electrode with the highest number of significantly decreased connections is CP4, the electrode overlapping with the area targeted by M1 TMS (Fig. 1 & 2).

**Figure 3:**
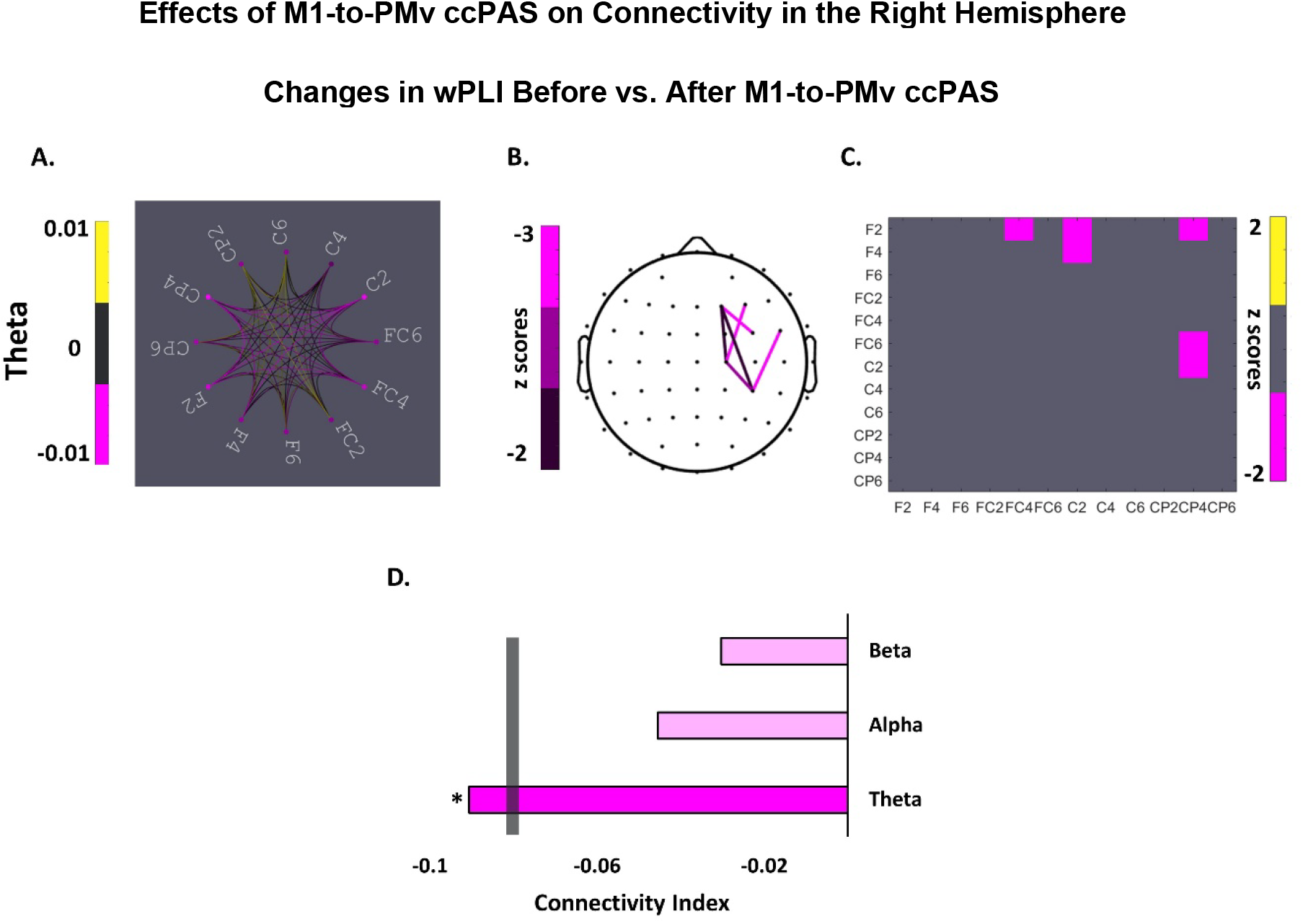
Changes in interregional coupling between PMv and M1 when contrasting activity recorded before and after M1-to-PMv ccPAS. (A) Representation of the connectivity strength measured by the weighted phase lag index (wPLI) across all electrodes included in the regions of interest for the theta band (B). Topographical representations of the electrodes showing significant decreased coupling in the theta band after M1-to-PMv ccPAS. (C) Connectivity matrix indicating significant interregional coupling decreases between the electrodes of interest. (D) Representation of the connectivity index for each of the frequency bands of interest; grey vertical bar shows the statistical threshold. Yellow and pink ink indicate increases or decreases in interregional coupling after ccPAS, respectively.

In addition, while in Group 1 there was a lack of change in interhemispheric connectivity (all CIs > CIthreshold), in Group 2 we observed a decrease in interhemispheric connectivity after the ccPAS protocol in the theta band (CItheta=-0,091 > CIthreshold), when computing the coherence between electrodes in the right and the left ROIs. Specifically, M1-to-PMv ccPAS led to a decrease of coherence between the electrodes in the vicinity of the right M1 TMS area (C2 and CP4) and their counterparts in the left hemisphere, electrodes C1 and CP3, as well as adjacent electrodes F5, FC3, FC5, C1, CP1, CP3 and CP5 (Fig. 4).

**Figure 4:**
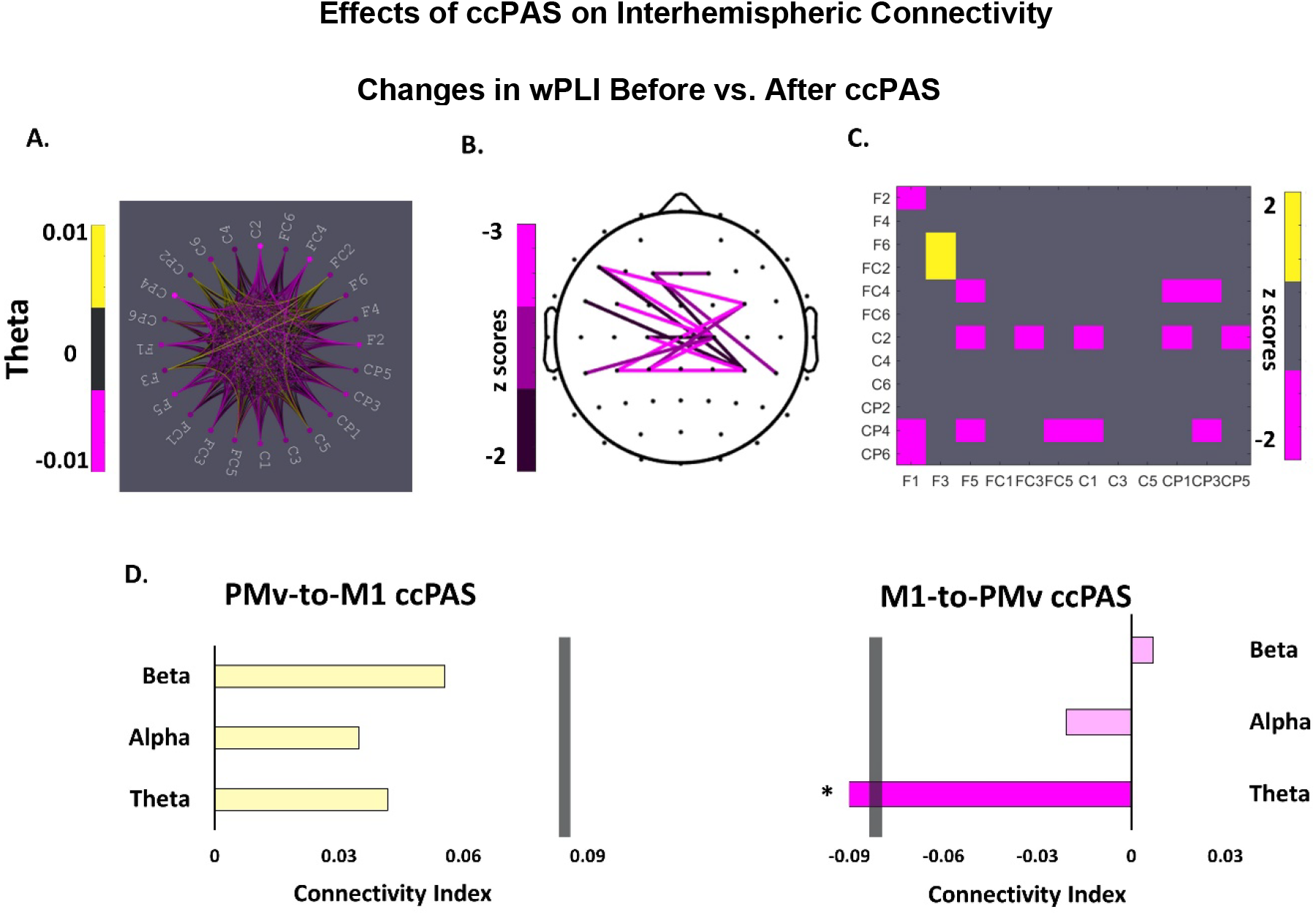
Changes in interregional coupling between PMv and M1 when contrasting activity recorded before and after ccPAS. (A) Representation of the connectivity strength measured by the weighted phase lag index (wPLI) across all electrodes included in the regions of interest in both hemispheres for the theta band in Group 2 (B) Topographical representations of the electrodes showing significant decreased coupling in the theta band after M1-to-PMv ccPAS. (C) Connectivity matrix indicating significant changes in interregional coupling between the electrodes of interest in Group 2. (D) Representation of the connectivity index for each of the frequency bands of interest in Group 1 (left) and Group 2 (right); grey vertical bar shows the statistical threshold. Yellow and pink ink indicate increases or decreases in interregional coupling after ccPAS, respectively.

Furthermore, we directly contrasted the connectivity changes between the two participant groups and across all frequency bands. We first computed the differences in wPLI by subtracting the connectivity indexes recorded at the baseline block from the connectivity indexes recorded in the expression block at the sites of interest – we called this the wPLI difference. We then directly contrasted the wPLI difference for each of the significant electrode cluster in each frequency bin (Theta, Alpha, Beta) between the two participant groups 1 and 2 (Fig. 5). The repeated measure analysis showed that the effects of the ccPAS on the resting-state connectivity are frequency- and cluster-specific across the two stimulation protocols (Frequency x Cluster x Group interaction: F (4, 136) = 3.285, p= .025, η_p_^2^ = .088).

**Figure 5:**
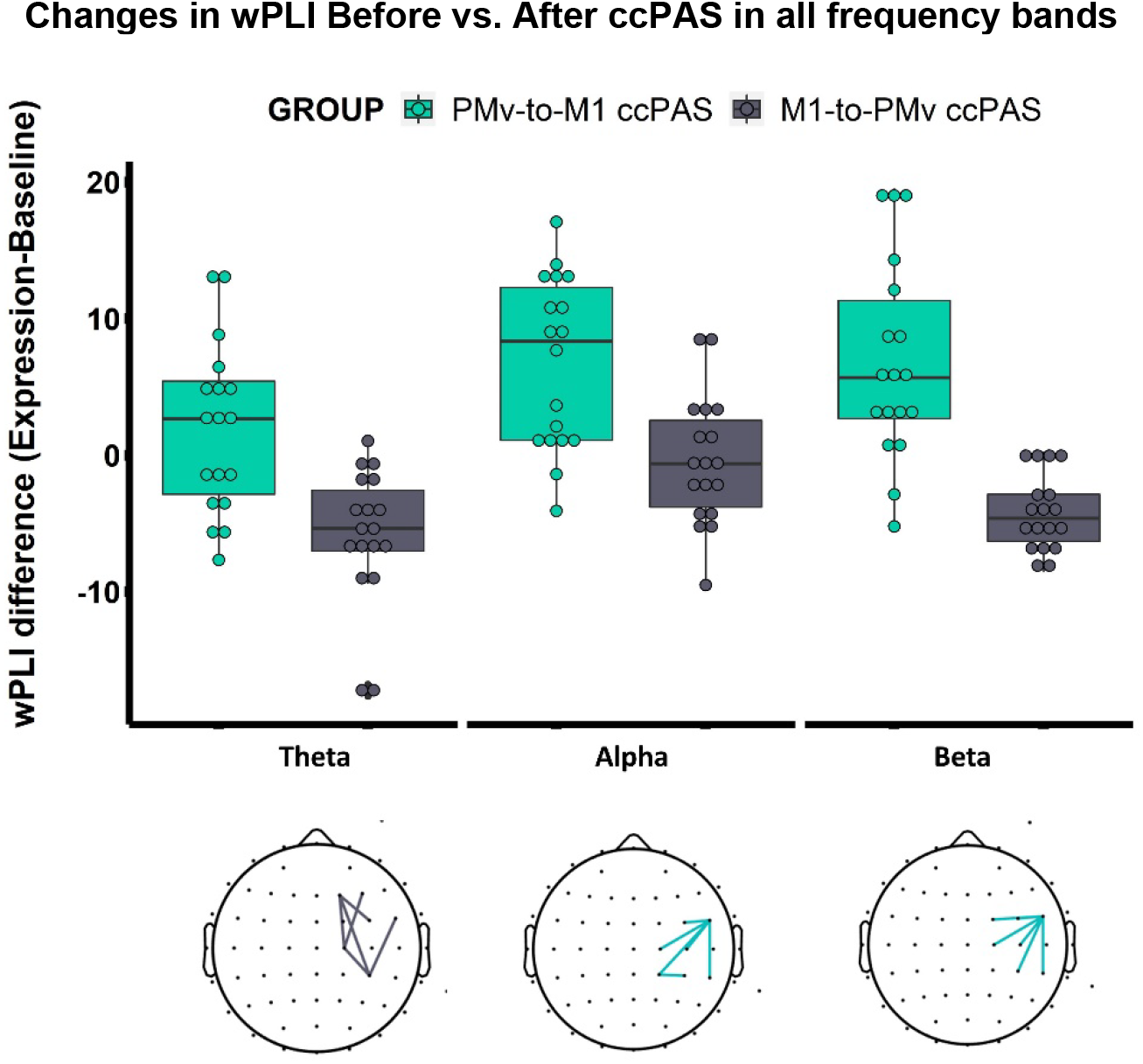
Changes in interregional connectivity for Group 1 (green) and Group 2 (grey) as measured by weighted phase lag index (wPLI) difference. The wPLI difference was computed by contrasting the wPLI recorded in the expression versus the baseline blocks, separately for the theta (left bars), alpha (middle bars), and beta (right bars) frequency bands. Each wPLI difference for each individual frequency was computed in the corresponding region of interest.

Specifically, the analysis confirmed that the increases and decreases in connectivity in the participant groups 1 and 2, respectively, occurred in the identified frequency-clusters in the matching frequency (See Fig. 5). This is, beta and alpha connectivity significantly increased in Group 1, as opposed to Group 2, when strengthening PMv-M1 connections; likewise, theta connectivity significantly decreased in Group 2 (*versus* Group 1) when decreasing PMv-M1 connections. Lastly, the results did not reveal connectivity changes within the left (non-stimulated) hemisphere that were significantly different between Groups 1 and 2 (all CIs > CI_threshold_) (Sup. Fig. 1), suggesting that the ccPAS effects were most apparent within the stimulated cortical sites and that the degree to which they were mirrored by parallel inter-areal coupling changes of the same scale in the unstimulated left hemisphere was limited.

Finally, we investigated the possible functional significance of these connectivity changes during resting state, and their relation to task-related changes. With this aim, we performed correlation analyses of ccPAS-induced connectivity changes during the resting-state and ccPAS-induced changes in time-frequency responses during a Go/No-go task performed by the same participants (both Groups 1 and 2) (Sel et al., 2021). Specifically, we used the wPLI as described above as a measure of changes when we examined connectivity strength between the EEG responses collected over PMv and M1 areas at rest by subtracting the signal recorded after ccPAS from the signal recorded before ccPAS.

Similarly, we subtracted changes in time-locked EEG power changes recorded while participants performed a motor task both making (‘going’) and stopping movements (Go/No-go task) after *versus* before the ccPAS protocol. We then examined the relationship between these two computed indices, one of intracortical communication at rest and one of task-related activity during motor control. We found a significant positive correlation between resting state and task-induced changes in both theta and beta oscillatory bands (beta band: h=1, r=0.385, CI = [0.015 0.701]; theta band: h=1, r=0.340, CI = [0.091 0.538]). Specifically, the greater the difference in resting-state connectivity before *versus* after ccPAS, the higher the effect of ccPAS on electrophysiological changes during the Go/No-go task (see Fig. 6). Overall, the malleability of the PMv-M1 connections that resulted from paired cortico-cortical stimulation that was visible at rest predicted increases and decreases in oscillatory activity during motor control.

**Figure 6.**
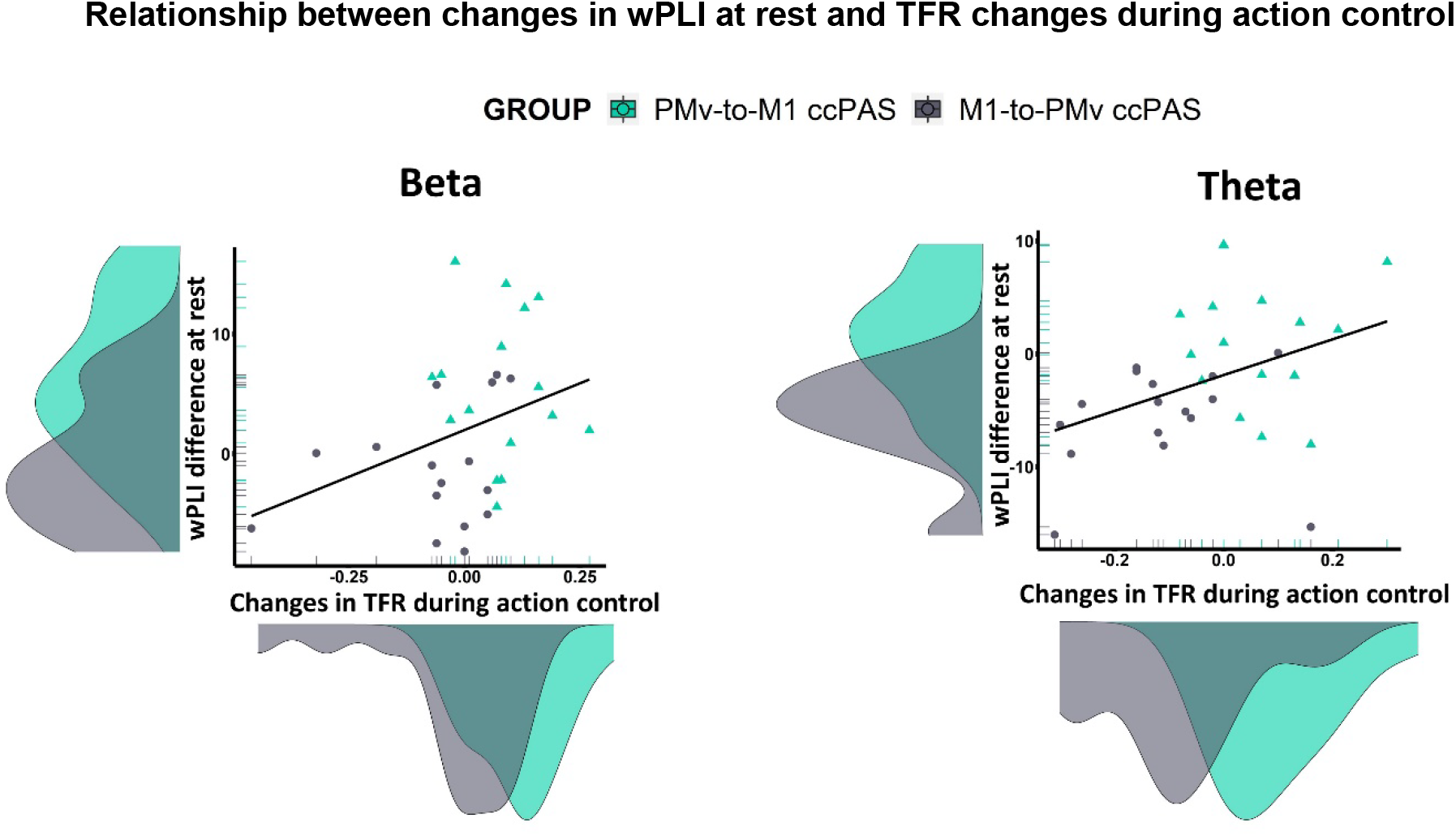
Correlation between interregional coherence changes and task-related changes in time frequency responses (TFR) before versus after ccPAS. Relationship between the changes in interregional coherence at rest – Y axis – and the changes in task-related oscillatory amplitude – X axis – in the beta (left) and theta (right) bands when contrasting activity recorded before and after ccPAS for both Groups 1 and 2 collectively. Green and grey ink indicate Group 1 and Group 2, respectively. Density distributions of the two variables are also presented along the corresponding axes.

## Discussion

There is wide interest in the possibility that brain oscillations are fundamental for communication between neuronal network elements and that their understanding might yield mechanistic insights into aspects of human cognition and behaviour (Schnitzler and Gross, 2005, Siegel et al., 2012, Fries, 2005). It has also been repeatedly suggested that the efficacy or strength of connections between neuronal groups influences the communication strength between brain regions (Fries, 2015, van Ede et al., 2018). Here we directly test this possibility in the human brain by using manipulations that have been established to either increase or decrease connectivity strength in a human cortico-cortical pathway, the route connecting PMv and M1. Depending on the inter-spike timing, ccPAS has been shown by several laboratories to result in either significant increases or decreases in functional coupling between PMv and M1 (Buch et al., 2011, Johnen et al., 2015, Casarotto et al., 2022, Chiappini et al., 2020, Turrini et al., 2022). Individual variation in functional connectivity between these regions has, in turn, been linked to individual variation in myelination of interconnecting pathways (Buch et al., 2011, Lazari et al., 2022b) and ccPAS-induced changes in functional connectivity are also associated with matched changes in myelination (Lazari et al., 2022a). We demonstrate that increasing and decreasing short-term synaptic efficacy increases and decreases interregional brain communication in the PMv-M1 pathway.

The repeated application of TMS pulses to PMv followed by M1, PMv-to-M1 ccPAS, evokes synchronous pre- and postsynaptic activity in the PMv-M1 pathway and increases the oscillatory communication in such a pathway. Specifically, beta and alpha oscillatory activity recorded over PMv and M1 regions became more phase-aligned following PMv-to-M1 ccPAS. This means that increasing connectivity in a motor control pathway of the human brain facilitates oscillatory communication between two anatomically connected motor control regions. Beta oscillatory activity is the dominant frequency band in interregional communication in the motor control circuit, particularly during inhibition and absence of movement (Picazio et al., 2014, Zhang et al., 2008, Ferreri et al., 2014), and the PMv-M1 pathway is a major cortical route by which the premotor cortex inhibits M1 motor-related activations at rest (Davare et al., 2008). Moreover, alpha oscillations are also linked to inhibitory control; increases in alpha activity over sensorimotor areas occurs in tandem with attenuated M1 motor-related activity and de-activation in premotor and motor cortical regions (Sauseng et al., 2009, Sauseng et al., 2013).

By contrast, reversing the order of stimulation, so that M1 TMS pulses are followed by PMv TMS pulses during the repeated stimulation protocols, leads to a decrease of interregional PMv-M1 coherence circumscribed to the theta band. It is unlikely that the reverse order M1-to-PMv stimulation protocol results in simultaneous pre- and post-synaptic activations in the PMv-M1 pathway; consequently, the synaptic efficacy and connectivity in the PMv-M1 pathway should either remain constant, or more likely, decrease (Buch et al., 2011, Johnen et al., 2015, Sel et al., 2021). According to the principles of Hebbian-like spike-timing dependent plasticity (Koch et al., 2013), the activation of presynaptic cells before postsynaptic cells leads to long-term potentiation whereas asynchronous activation of the neurons or activation of postsynaptic neurons before presynaptic neurons induces long-term depression.

Together, the findings demonstrate that it is possible to selectively modulate functional connectivity in an important motor control circuit in the human brain by recurrent stimulation of the pathway. They also suggest that transmission of causal influences in the cortical pathway connecting the PMv and M1 are instantiated in separate channels of communications tunned to certain frequencies. Specifically, the faster beta and alpha rhythms and the slower theta rhythm orchestrate distinct aspects of top-down control over motor cortex.

Different cortical rhythms in the beta, alpha and theta range are associated with distinct functional roles in executive control (Tsujimoto et al., 2010, Tsujimoto et al., 2006, Yamanaka and Yamamoto, 2010, Harper et al., 2014, Helfrich et al., 2018, Helfrich et al., 2019, Picazio et al., 2014). These oscillatory response patterns are instrumental for conveying information from the premotor cortex to M1 cortex during movement selection and cessation (Picazio et al., 2014, Schnitzler and Gross, 2005, Siegel et al., 2012). For example, augmentation of beta power in prefrontal regions is associated with increased motor control and precision in movement timing; in turn, these increases in beta power are known to be driven by increases in alpha phase coupling between frontal and central regions reflecting integration of information across premotor and motor areas during top-down executive control (Kononowicz et al., 2020, Grabot et al., 2019). In addition, increases in theta phase coupling between frontal and central regions within the same hemisphere are observed during action inhibition and action reprogramming (Shibata et al., 1997, Shibata et al., 1998). Here we observe that individuals that exhibited greater increases in fronto-central beta phase coupling after PMv-to-M1 ccPAS also showed selective enhancement of movement-induced beta power at the time of movement completion before *versus* after ccPAS. In a similar vein, those individuals exhibiting the greatest decreases in interregional phase coupling in the theta band after undergoing ccPAS in the reversed M1-to-PMv order also presented the largest reductions in theta power during action inhibition following reversed order M1-to-PMv ccPAS which weakens the PMv-M1 pathway (Sel et al., 2021). Collectively, these results indicate that the neuronal architecture supporting the PMv-M1 network has fundamental resonant properties in different frequency bands and that even, in the absence of any motor task, different manipulations of PMv-M1 pathway strength affect specific communication channels. However, the functional impact of changes in the different channels may be most apparent during different aspects of motor behaviour such as action execution and inhibition. The results also suggest that it is possible to anticipate the impact that a given manipulation will have on an aspect of behaviour given prior knowledge of 1) the impact of a PMv-M1 manipulation at rest on different frequency channels; 2) the association between the frequency channels and behaviour.

Increases in beta and alpha interregional coherence changes were circumscribed to the sites of stimulation over right PMv and right M1 and adjacent cortical areas. Beta and alpha phase coupling was observed predominantly at the right lateral frontal sensor - particularly electrode FC6 – located over PMv and adjacent inferior frontal cortex. The phase of activity in this electrode became more aligned to the phase of the beta and alpha activity in electrodes placed over the right M1 and neighbouring sensorimotor cortices, following PMv-to-M1 ccPAS. Conversely, reversed order M1-to-PMv ccPAS led to a decrease in phase alignment in the theta band distributed across both hemispheres; activity in sensorimotor electrodes became more misaligned with activity recorded over medial and lateral frontocentral sites. Cortical activity within the beta, alpha and theta bands occur in medial and lateral frontal areas that interact with PMv during action inhibition (Tsujimoto et al., 2006, Tsujimoto et al., 2010, Aron et al., 2014, Aron et al., 2007, Neubert et al., 2010). These areas include the pre-SMA in the dorsal frontomedial cortex, inferior frontal cortex anterior to PMv itself, and M1 (Aron et al., 2014, Aron et al., 2007, Neubert et al., 2010). In line with this observation, analysis of BOLD coupling at rest in areas that interact with PMv and M1 has established that augmented pathway efficacy following PMv-to-M1 ccPAS is mirrored in increased interregional functional connectivity within the PMv-M1 pathway, as well as between PMv and the broader frontal-parietal motor association network (Johnen et al., 2015). The current findings complement previous evidence of increased functional connectivity at rest after ccPAS (Veniero et al., 2013), and of selective enhancement of specific pathways beyond just PMv-M1 itself (Chiappini et al., 2018, Santarnecchi et al., 2018).

It is possible that the oscillatory changes observed in the beta, alpha and theta bands are linked to changes in the phase alignment occurring primarily in prefrontal regions which then spread to PMv and M1 to produce the observed increases and decreases of cortico-cortical coupling between PMv and M1. However, this possibility is unlikely because any changes in phase alignment resulting from volume conductance, or changes occurring at a distant site, have been controlled by computing the weighted phase lag indexes, an unbiased estimator with low sensitivity to volume-conduction and uncorrelated noise sources (Vinck et al., 2011).

Overall, our results indicate that functional connectivity as indexed by interregional synchronisation of rhythmic activity can reflect short-term increases in synaptic efficacy and neural wiring. The synchronisation of activity across different sets of neuronal groups is only possible if they share a common neurophysiological substrate (Fries, 2015, Fries, 2005, Salinas and Sejnowski, 2001); when this physical subtract becomes more tightly wired as a result of sensory learning (Garrido et al., 2009) or motor training (Ackerley et al., 2011), the brain frequency activity governing communication between the neuronal network elements align, oscillating with similar rhythmic patterns (Plewnia et al., 2008). On the contrary, reductions in synaptic efficacy observed, for example, after medication in Parkinson’s disease, are mirrored in suppression of beta phase alignment within deeper structures of the motor control network (van Wijk et al., 2018). Our results are also consistent with previous investigations demonstrating, first, that inter-individual variation in myelination in PMv-related anatomical pathways is related to inter-individual variation in the impact that the induction of activity in PMv (with a TMS pulse) has on activity in M1 (Lazari et al., 2022b), and second that ccPAS induces measurable changes in the white matter track (Lazari et al., 2022a).

## Conclusions

In summary, these results illustrate a mechanistic link binding synaptic efficacy in short-range connections to their functional role as conveyors of information communication through brain oscillatory transmissions. They highlight, for the first time, that interregional communication frequencies in the human PMv-M1 pathway can be manipulated leading to increases or decreases in frequency phase alignment of the oscillatory architecture supporting action control. The selective frequency-specific patterns of phase coupling change found after different types of ccPAS reflect spectral fingerprints of augmentation *versus* reduction of top-down PMv influence over M1. Taken together, these results are consistent with Hebbian-like spike-timing dependent long-term potentiation and depression (Koch et al., 2013) and with hierarchical models of action control in which top-down executive control occurs in tandem with phase coupling with specific resonant properties in the beta, alpha and theta frequency ranges (Helfrich et al., 2018, Helfrich et al., 2019, Sauseng et al., 2009).

## Data availability

Anonymised human brain data and analysis scripts have been deposited in the Open Science Framework repository (https://osf.io/4k9jn/?view_only=01cd3db80dff404cbb47c56904f43b2e).

## Acknowledgments

Funders: Bial Foundation (Grant 44/16) to AS, and the Wellcome Trust (221794/Z/20/Z) and John Templeton Foundation Prime Award (15464/ Subaward Ref. SC14) to MFSR. We acknowledge the assistance of Nadescha Trudel, Raluca David, Katarina Angerer during data collection.

## Author Contributions

Conceived and designed the experiments: AS MFSR. Performed the experiments: AS. Analyzed the data: JT AS VR MFSR. Wrote the paper: AS JT VR MFSR.

## References

Ackerley, S. J., Stinear, C. M. & Byblow, W. D. 2011. Promoting use-dependent plasticity with externally-paced training. Clinical Neurophysiology, 122, 2462–2468.

Alekseichuk, I., Turi, Z., Amador de Lara, G., Antal, A. & Paulus, W. 2016. Spatial Working Memory in Humans Depends on Theta and High Gamma Synchronization in the Prefrontal Cortex. Current Biology, 26, 1513–1521.

Aron, A. R., Behrens, T. E., Smith, S., Frank, M. J. & Poldrack, R. A. 2007. Triangulating a cognitive control network using diffusion-weighted magnetic resonance imaging (MRI) and functional MRI. J Neurosci, 27, 3743–52.

Aron, A. R., Robbins, T. W. & Poldrack, R. A. 2014. Inhibition and the right inferior frontal cortex: one decade on. Trends in cognitive sciences, 18, 177–185.

Buch, E. R., Johnen, V. M., Nelissen, N., O’shea, J. & Rushworth, M. F. 2011. Noninvasive associative plasticity induction in a corticocortical pathway of the human brain. J Neurosci, 31, 17669–79.

Buch, E. R., Mars, R. B., Boorman, E. D. & Rushworth, M. F. 2010. A network centered on ventral premotor cortex exerts both facilitatory and inhibitory control over primary motor cortex during action reprogramming. J Neurosci, 30, 1395–401.

Casarotto, A., Dolfini, E., Cardellicchio, P., Fadiga, L., D’ausilio, A. & Koch, G. 2022. Mechanisms of Hebbian-like plasticity in the ventral premotor–primary motor network. The Journal of Physiology.

Cerri, G., Shimazu, H., Maier, M. A. & Lemon, R. N. 2003. Facilitation from ventral premotor cortex of primary motor cortex outputs to macaque hand muscles. J Neurophysiol, 90, 832–42.

Chiappini, E., Borgomaneri, S., Marangon, M., Turrini, S., Romei, V. & Avenanti, A. 2020. Driving associative plasticity in premotor-motor connections through a novel paired associative stimulation based on long-latency cortico-cortical interactions. Brain Stimulation: Basic, Translational, and Clinical Research in Neuromodulation, 13, 1461–1463.

Chiappini, E., Sel, A., Hibbard, P. B., Avenanti, A. & Romei, V. 2022. Increasing interhemispheric connectivity between human visual motion areas uncovers asymmetric sensitivity to horizontal motion. Current Biology.

Chiappini, E., Silvanto, J., Hibbard, P. B., Avenanti, A. & Romei, V. 2018. Strengthening functionally specific neural pathways with transcranial brain stimulation. Current Biology, 28, R735–R736.

Cohen, M. X. 2014. Analyzing neural time series data: theory and practice, MIT press.

Cohen, M. X. 2015. Effects of time lag and frequency matching on phase-based connectivity. Journal of Neuroscience Methods, 250, 137–146.

Croxson, P. L., Johansen-Berg, H., Behrens, T. E. J., Robson, M. D., Pinsk, M. A., Gross, C. G., Richter, W., Richter, M. C., Kastner, S. & Rushworth, M. F. S. 2005. Quantitative Investigation of Connections of the Prefrontal Cortex in the Human and Macaque using Probabilistic Diffusion Tractography. The Journal of Neuroscience, 25, 8854–8866.

Davare, M., Kraskov, A., Rothwell, J. C. & Lemon, R. N. 2011. Interactions between areas of the cortical grasping network. Current Opinion in Neurobiology, 21, 565–570.

Davare, M., Lemon, R. & Olivier, E. 2008. Selective modulation of interactions between ventral premotor cortex and primary motor cortex during precision grasping in humans. The Journal of Physiology, 586, 2735–2742.

Davare, M., Montague, K., Olivier, E., Rothwell, J. C. & Lemon, R. N. 2009. Ventral premotor to primary motor cortical interactions during object-driven grasp in humans. Cortex, 45, 1050–1057.

Davare, M., Rothwell, J. C. & Lemon, R. N. 2010. Causal connectivity between the human anterior intraparietal area and premotor cortex during grasp. Current Biology, 20, 176–181.

Dum, R. P. & Strick, P. L. 2005. Frontal lobe inputs to the digit representations of the motor areas on the lateral surface of the hemisphere. Journal of Neuroscience, 25, 1375–1386.

Ferreri, F., Vecchio, F., Ponzo, D., Pasqualetti, P. & Rossini, P. M. 2014. Time-varying coupling of EEG oscillations predicts excitability fluctuations in the primary motor cortex as reflected by motor evoked potentials amplitude: An EEG-TMS study. Human Brain Mapping, 35, 1969–1980.

Fiori, F., Chiappini, E. & Avenanti, A. 2018. Enhanced action performance following TMS manipulation of associative plasticity in ventral premotor-motor pathway. NeuroImage, 183, 847–858.

Fries, P. 2005. A mechanism for cognitive dynamics: neuronal communication through neuronal coherence. Trends in cognitive sciences, 9, 474–480.

Fries, P. 2015. Rhythms for cognition: communication through coherence. Neuron, 88, 220–235.

Garrido, M. I., Kilner, J. M., Kiebel, S. J., Stephan, K. E., Baldeweg, T. & Friston, K. J. 2009. Repetition suppression and plasticity in the human brain. Neuroimage, 48, 269–79.

Grabot, L., Kononowicz, T. W., La Tour, T. D., Gramfort, A., Doyère, V. & Van Wassenhove, V. 2019. The strength of alpha–beta oscillatory coupling predicts motor timing precision. Journal of Neuroscience, 39, 3277–3291.

Harper, J., Malone, S. M. & Bernat, E. M. 2014. Theta and delta band activity explain N2 and P3 ERP component activity in a go/no-go task. Clinical Neurophysiology, 125, 124–132.

Helfrich, R. F., Breska, A. & Knight, R. T. 2019. Neural entrainment and network resonance in support of top-down guided attention. Curr Opin Psychol, 29, 82–89.

Helfrich, R. F., Fiebelkorn, I. C., Szczepanski, S. M., Lin, J. J., Parvizi, J., Knight, R. T. & Kastner, S. 2018. Neural Mechanisms of Sustained Attention Are Rhythmic. Neuron, 99, 854–865 e5.

Huang, Y. Z., Lu, M. K., Antal, A., Classen, J., Nitsche, M., Ziemann, U., Ridding, M., Hamada, M., Ugawa, Y., Jaberzadeh, S., Suppa, A., Paulus, W. & Rothwell, J. 2017. Plasticity induced by non-invasive transcranial brain stimulation: A position paper. Clin Neurophysiol, 128, 2318–2329.

Hughes, L. E., Rittman, T., Robbins, T. W. & Rowe, J. B. 2018. Reorganization of cortical oscillatory dynamics underlying disinhibition in frontotemporal dementia. Brain, 141, 2486–2499.

Johnen, V. M., Neubert, F. X., Buch, E. R., Verhagen, L., O’reilly, J. X., Mars, R. B. & Rushworth, M. F. 2015. Causal manipulation of functional connectivity in a specific neural pathway during behaviour and at rest. Elife, 4.

Koch, G., Ponzo, V., Di Lorenzo, F., Caltagirone, C. & Veniero, D. 2013. Hebbian and Anti-Hebbian Spike-Timing-Dependent Plasticity of Human Cortico-Cortical Connections. The Journal of Neuroscience, 33, 9725–9733.

Kononowicz, T. W., Sander, T., Van Rijn, H. & Van Wassenhove, V. 2020. Precision Timing with α-β Oscillatory Coupling: Stopwatch or Motor Control? J Cogn Neurosci, 32, 1624–1636.

Lazari, A., Salvan, P., Cottaar, M., Papp, D., Rushworth, M. F. S. & Johansen-Berg, H. 2022a. Hebbian activity-dependent plasticity in white matter. Cell Rep, 39, 110951.

Lazari, A., Salvan, P., Verhagen, L., Cottaar, M., Papp, D., Van Der Werf, O. J., Gavine, B., Kolasinski, J., Webster, M. & Stagg, C. J. 2022b. A macroscopic link between interhemispheric tract myelination and cortico-cortical interactions during action reprogramming. Nature communications, 13, 1–12.

Liebrand, M., Kristek, J., Tzvi, E. & Krämer, U. M. 2018. Ready for change: Oscillatory mechanisms of proactive motor control. PloS one, 13, e0196855.

Mayka, M. A., Corcos, D. M., Leurgans, S. E. & Vaillancourt, D. E. 2006. Three-dimensional locations and boundaries of motor and premotor cortices as defined by functional brain imaging: A meta-analysis. NeuroImage, 31, 1453–1474.

Neubert, F.-X., Mars, ROGIER B., Thomas, ADAM G., Sallet, J. & Rushworth, MATTHEW F. S. 2014. Comparison of Human Ventral Frontal Cortex Areas for Cognitive Control and Language with Areas in Monkey Frontal Cortex. Neuron, 81, 700–713.

Neubert, F. X., Mars, R. B., Buch, E. R., Olivier, E. & Rushworth, M. F. 2010. Cortical and subcortical interactions during action reprogramming and their related white matter pathways. Proc Natl Acad Sci U S A, 107, 13240–5.

Oldfield, R. C. 1971. The assessment and analysis of handedness: The Edinburgh inventory. Neuropsychologia, 9, 97–113.

Oostenveld, R., Fries, P., Maris, E. & Schoffelen, J.-M. 2011. FieldTrip: Open Source Software for Advanced Analysis of MEG, EEG, and Invasive Electrophysiological Data. Computational Intelligence and Neuroscience, 2011, 156869.

Pernet, C., Wilcox, R. & Rousselet, G. 2013. Robust Correlation Analyses: False Positive and Power Validation Using a New Open Source Matlab Toolbox. Frontiers in Psychology, 3.

Picazio, S., Veniero, D., Ponzo, V., Caltagirone, C., Gross, J., Thut, G. & Koch, G. 2014. Prefrontal Control over Motor Cortex Cycles at Beta Frequency during Movement Inhibition. Current Biology, 24, 2940–2945.

Plewnia, C., Rilk, A. J., Soekadar, S. R., Arfeller, C., Huber, H. S., Sauseng, P., Hummel, F. & Gerloff, C. 2008. Enhancement of long-range EEG coherence by synchronous bifocal transcranial magnetic stimulation. Eur J Neurosci, 27, 1577–83.

Prabhu, G., Shimazu, H., Cerri, G., Brochier, T., Spinks, R. L., Maier, M. A. & Lemon, R. N. 2009a. Modulation of primary motor cortex outputs from ventral premotor cortex during visually guided grasp in the macaque monkey. The Journal of physiology, 587, 1057–1069.

Prabhu, G., Shimazu, H., Cerri, G., Brochier, T., Spinks, R. L., Maier, M. A. & Lemon, R. N. 2009b. Modulation of primary motor cortex outputs from ventral premotor cortex during visually guided grasp in the macaque monkey. J Physiol, 587, 1057–69.

Romei, V., Chiappini, E., Hibbard, PAUL B. & Avenanti, A. 2016. Empowering Reentrant Projections from V5 to V1 Boosts Sensitivity to Motion. Current Biology, 26, 2155–2160.

Romero, M. C., Davare, M., Armendariz, M. & Janssen, P. 2019. Neural effects of transcranial magnetic stimulation at the single-cell level. Nat Commun, 10, 2642.

Rossini, P. M., Barker, A., Berardelli, A., Caramia, M., Caruso, G., Cracco, R., Dimitrijeviš, M., Hallett, M., Katayama, Y. & Lücking, C. 1994. Non-invasive electrical and magnetic stimulation of the brain, spinal cord and roots: basic principles and procedures for routine clinical application. Report of an IFCN committee. Electroencephalography and clinical neurophysiology, 91, 79–92.

Salinas, E. & Sejnowski, T. J. 2001. Correlated neuronal activity and the flow of neural information. Nature reviews neuroscience, 2, 539–550.

Santarnecchi, E., Momi, D., Sprugnoli, G., Neri, F., Pascual-Leone, A., Rossi, A. & Rossi, S. 2018. Modulation of network-to-network connectivity via spike-timing-dependent noninvasive brain stimulation. Human Brain Mapping, 39, 4870–4883.

Sauseng, P., Gerloff, C. & Hummel, F. C. 2013. Two brakes are better than one: the neural bases of inhibitory control of motor memory traces. Neuroimage, 65, 52–58.

Sauseng, P., Klimesch, W., Gerloff, C. & Hummel, F. C. 2009. Spontaneous locally restricted EEG alpha activity determines cortical excitability in the motor cortex. Neuropsychologia, 47, 284–288.

Schnitzler, A. & Gross, J. 2005. Normal and pathological oscillatory communication in the brain. Nature reviews neuroscience, 6, 285–296.

Sel, A., Verhagen, L., Angerer, K., David, R., Klein-flügge, M. C. & Rushworth, M. F. S. 2021. Increasing and decreasing interregional brain coupling increases and decreases oscillatory activity in the human brain. Proceedings of the National Academy of Sciences, 118, e2100652118.

Shibata, T., Shimoyama, I., Ito, T., Abla, D., Iwasa, H., Koseki, K., Yamanouchi, N., Sato, T. & Nakajima, Y. 1997. The time course of interhemispheric EEG coherence during a GO/NO-GO task in humans. Neuroscience letters, 233, 117–120.

Shibata, T., Shimoyama, I., Ito, T., Abla, D., Iwasa, H., Koseki, K., Yamanouchi, N., Sato, T. & Nakajima, Y. 1998. The synchronization between brain areas under motor inhibition process in humans estimated by event-related EEG coherence. Neuroscience research, 31, 265–271.

Shimazu, H., Maier, M. A., Cerri, G., Kirkwood, P. A. & Lemon, R. N. 2004. Macaque Ventral Premotor Cortex Exerts Powerful Facilitation of Motor Cortex Outputs to Upper Limb Motoneurons. J Neurosci, 24, 1200–1211.

Siegel, M., Donner, T. H. & Engel, A. K. 2012. Spectral fingerprints of large-scale neuronal interactions. Nature Reviews Neuroscience, 13, 121–134.

Tokuno, H. & Nambu, A. 2000. Organization of nonprimary motor cortical inputs on pyramidal and nonpyramidal tract neurons of primary motor cortex: an electrophysiological study in the macaque monkey. Cerebral cortex, 10, 58–68.

Tsujimoto, T., Shimazu, H. & Isomura, Y. 2006. Direct recording of theta oscillations in primate prefrontal and anterior cingulate cortices. J Neurophysiol, 95, 2987–3000.

Tsujimoto, T., Shimazu, H., Isomura, Y. & Sasaki, K. 2010. Theta oscillations in primate prefrontal and anterior cingulate cortices in forewarned reaction time tasks. J Neurophysiol, 103, 827–43.

Turrini, S., Fiori, F., Chiappini, E., Santarnecchi, E., Romei, V. & Avenanti, A. 2022. Gradual enhancement of corticomotor excitability during cortico-cortical paired associative stimulation. Scientific Reports, 12, 1–8.

Van Ede, F., Quinn, A. J., Woolrich, M. W. & Nobre, A. C. 2018. Neural oscillations: sustained rhythms or transient burst-events? Trends in neurosciences, 41, 415–417.

Van Wijk, B. C., Cagnan, H., Litvak, V., Kühn, A. A. & Friston, K. J. 2018. Generic dynamic causal modelling: An illustrative application to Parkinson’s disease. NeuroImage, 181, 818–830.

Veniero, D., Ponzo, V. & Koch, G. 2013. Paired Associative Stimulation Enforces the Communication between Interconnected Areas. The Journal of Neuroscience, 33, 13773–13783.

Vinck, M., Oostenveld, R., Van Wingerden, M., Battaglia, F. & Pennartz, C. M. A. 2011. An improved index of phase-synchronization for electrophysiological data in the presence of volume-conduction, noise and sample-size bias. NeuroImage, 55, 1548–1565.

Wang, X.-J. 2010. Neurophysiological and computational principles of cortical rhythms in cognition. Physiological reviews, 90, 1195–1268.

Yamanaka, K. & Yamamoto, Y. 2010. Single-trial EEG Power and Phase Dynamics Associated with Voluntary Response Inhibition. Journal of Cognitive Neuroscience, 22, 714–727.

Zhang, Y., Chen, Y., Bressler, S. L. & Ding, M. 2008. Response preparation and inhibition: The role of the cortical sensorimotor beta rhythm. Neuroscience, 156, 238–246.

