## Supplementary Figure 1 for "Strengthening connectivity between premotor and motor cortex increases inter-areal communication in the human brain"

**Effects of ccPAS on Connectivity in the Left Hemisphere**

**Changes in wPLI Before vs. After ccPAS**

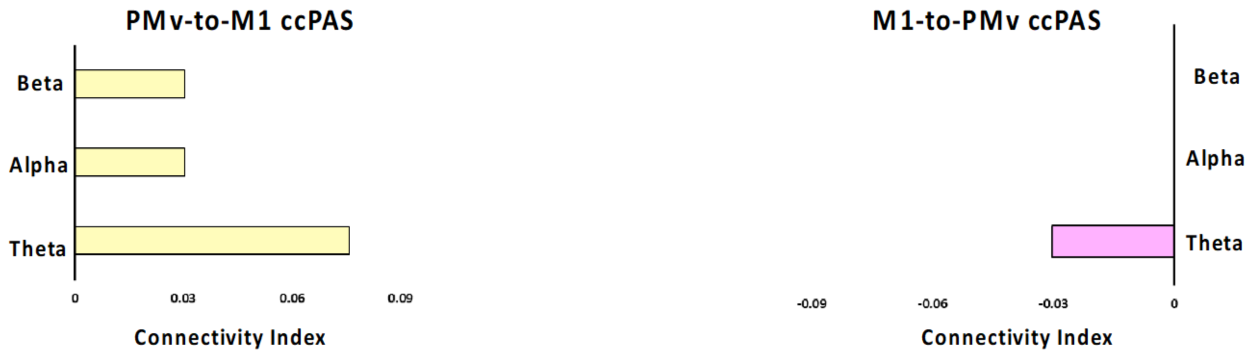


***Supplementary Figure 1**: **Representation of the connectivity index for each of the frequency bands of interest in Group 1 (left) and Group 2 (right) in the left hemisphere.** Connectivity results found when recording changes in interregional coupling measured by the weighted phase lag index (wPLI) in the left (non-stimulated) hemisphere; grey vertical bar shows the statistical threshold.*
